# Whole exome precision oncology targeting synthetic lethal vulnerabilities across the tumor transcriptome

**DOI:** 10.1101/2020.02.16.951699

**Authors:** Joo Sang Lee, Nishanth Ulhas Nair, Lesley Chapman, Sanju Sinha, Kun Wang, Houngui Cha, Eitan Rubin, Raanan Berger, Vladimir Lazar, Razelle Kurzrock, Mark R. Gilbert, Sridhar Hannenhalli, Se-Hoon Lee, Kenneth Aldape, Eytan Ruppin

**Affiliations:** Cancer Data Science Lab, Center for Cancer Research, National Cancer Institute, National Institutes of Health, Bethesda MD, USA; Samsung Medical Center, Sungkyunkwan University School of Medicine, Suwon, Republic of Korea; Ben-Gurion University of the Negev, Beersheva, Israel; Chaim Sheba Medical Center, Tel Hashomer, Israel; Worldwide Innovative Network (WIN) Association-WIN Consortium, Villejuif, France; University of California San Diego, Moore Cancer Center, San Diego, CA, USA; Neuro-Oncology Branch, Center for Cancer Research, National Cancer Institute, National Institutes of Health, Bethesda MD, USA; Laboratory of Pathology, Center for Cancer Research, National Cancer Institute, National Institutes of Health, Bethesda MD, USA

## Abstract

Precision oncology has made significant advances in the last few years, mainly by targeting actionable mutations in cancer driver genes. However, the proportion of patients whose tumors can be targeted therapeutically remains limited. Recent studies have begun to explore the benefit of analyzing tumor transcriptomics data to guide patient treatment, raising the need for new approaches for systematically accomplishing that. Here we show that computationally derived genetic interactions can successfully predict patient response. Assembling a broad repertoire of 32 datasets spanning more than 1,500 patients and including both tumor transcriptomics and response data, we predicted the response in 17 out of 21 targeted and 8 out of 11 checkpoint therapy datasets across 8 different cancer types with considerable accuracy, without ever training on these datasets. Analyzing the recently published multi-arm WINTHER trial, we show that the fraction of patients benefitting from transcriptomic-based treatments could potentially be markedly increased from 15% to about 85% by targeting synthetic lethal vulnerabilities in their tumors. In summary, this is the first computational approach to obtain considerable predictive performance across many different targeted and immunotherapy datasets, providing a promising new way for guiding cancer treatment based on the tumor transcriptomics of cancer patients.

## Introduction

There have been significant advances in precision oncology, with an increasing adoption of sequencing tests that identify targetable mutations in cancer driver genes. Aiming to complement these efforts by considering genome-wide tumor alterations at additional “-omics” layers, recent studies have begun to explore the utilization of transcriptomics data to guide cancer patients’ treatment^1-4^. These studies have reported encouraging results, testifying to the potential of such approaches to complement mutation panels and increase the likelihood that patients will benefit from genomics-guided, precision treatments. However, current approaches have still been of a heuristic exploratory nature, raising the need for developing and testing new systematic approaches for utilizing tumor transcriptomics data.

Here we present a novel precision oncology framework aimed at stratifying patients to existing cancer targeted and immunotherapy drugs, based on gene expression data from their tumors. Our main goal is to extend the scope of current approaches from a few hundred driver genes to genomic and transcriptomic alterations occurring across the *whole exome*. Unlike the recent tumor transcriptome-based precision oncology approaches resorting mainly to the expression of drug targets, our approach is based on identifying and utilizing the broader scope of *genetic interactions* (GI) of the drug target genes. We focus on two major types that are highly relevant to cancer therapies: (1) *Synthetic lethal (SL) interactions*, which describe the relationship between two genes whose concomitant inactivation is required to reduce the viability of the cell (e.g., an SL interaction that is widely used in the clinic is of PARP inhibitors on the background of disrupted DNA repair)^5^. (2) *Synthetic rescue (SR) interactions*, which denote a type of genetic interaction where a change in the activity of one gene reduces the cell’s fitness but an alteration of another gene (termed its SR partner) rescues cell viability (e.g., the rescue of Myc alterations by BCL2 activation in lymphomas^6^). When a gene is targeted by a small molecule inhibitor or an antibody, the tumor may respond by changing the activity of its rescuer gene(s), conferring resistance to therapies.

To identify the SL and SR partners of cancer drugs, we leverage two recently published novel computational pipelines, ISLE^7^ and INCISOR^8^, respectively, which extract genetic dependencies that are supported by multiple layers of omics data, including *in vitro* functional screens, patient tumor DNA and RNA sequencing data, and phylogenic similarity across different species. The ISLE pipeline has recently successfully identified a Gq-driver mutation as marker for FAK inhibitor treatment in uveal melanoma^9^ and a synergistic combination for treating melanoma and pancreatic tumors with Asparaginase and MAPK inhibitors^10^. Similarly, INCISOR analysis has identified SR interactions that mediate resistance to checkpoint therapies in melanoma^8^. Here, we set out to evaluate comprehensively for the first time the clinical utility of the computationally inferred GIs as tumor transcriptome-based companion diagnostics for stratifying cancer patients to a host of targeted and immunotherapy drugs in new, ‘unseen’ and independent, patients’ treatment cohorts.

## Results

### Overview of the analysis

We collected cancer patients’ pre-treatment transcriptomics profiles together with therapy response information from numerous publicly available databases, surveying Gene Expression Omnibus (GEO), ArrayExpress and the literature, and a new unpublished cohort of anti-PD1 treatment in lung adenocarcinoma. Overall, we found 32 such datasets that include both transcriptomics and clinical response data, spanning 21 targeted therapies and 11 immunotherapy datasets across 10 different cancer types.

For each drug whose response we aim to predict, we first applied the ISLE^7^ and INCISOR^8^ pipelines to identify the *clinically relevant* pan-cancer GIs (the interactions found to be shared across many cancer types) of its target genes^11^. Here we briefly describe the process for identifying and generating predictions based on SLs, and the process for using SRs is analogous (see **Figure 1A** and Methods): (A) <u>SL inference</u>: For each drug we compile a list of initial candidate SL pairs of its targets by analyzing large-scale *in vitro* functional screens performed with RNAi, CRISPR/Cas9, or pharmacological inhibition in DepMap^12,13^. Among these candidate SL pairs, we then select those that are more likely to be clinically relevant by analyzing the TCGA data. Finally, among the remaining candidate pairs we select those pairs that are supported by a phylogenetic profiling analysis^7^. The top 25 SL partners that pass all three filters are designated as the SL partners of that drug. (B) <u>SL-based response prediction</u>: the identified SL partners of the drug are then used to predict a given patient’s response to a given treatment based on her/his tumor’s gene expression. This is based on the notion that a drug will kill a tumor more effectively when its SL partners are down-regulated, because when the drug inhibits its targets more SL interactions will become jointly down-regulated and hence ‘activated’ (**Figure 1B**). To quantify the extent of such predicted lethality, we assign an *SL-score* denoting the fraction of down-regulated SL partners of a drug in a given tumor (Methods). The larger the fraction of SL partners being down-regulated, the higher the SL-score and the more likely the patient is to respond to the given therapy. Analogously for the case of immunotherapy, we use an SR-score for predicting treatment outcomes, which quantifies the differences in the fraction of up or down regulated predicted SR partner genes based on the patient’s tumor transcriptomics, and hence the likelihood of resistance to the given therapies (Methods).

**Figure 1.**
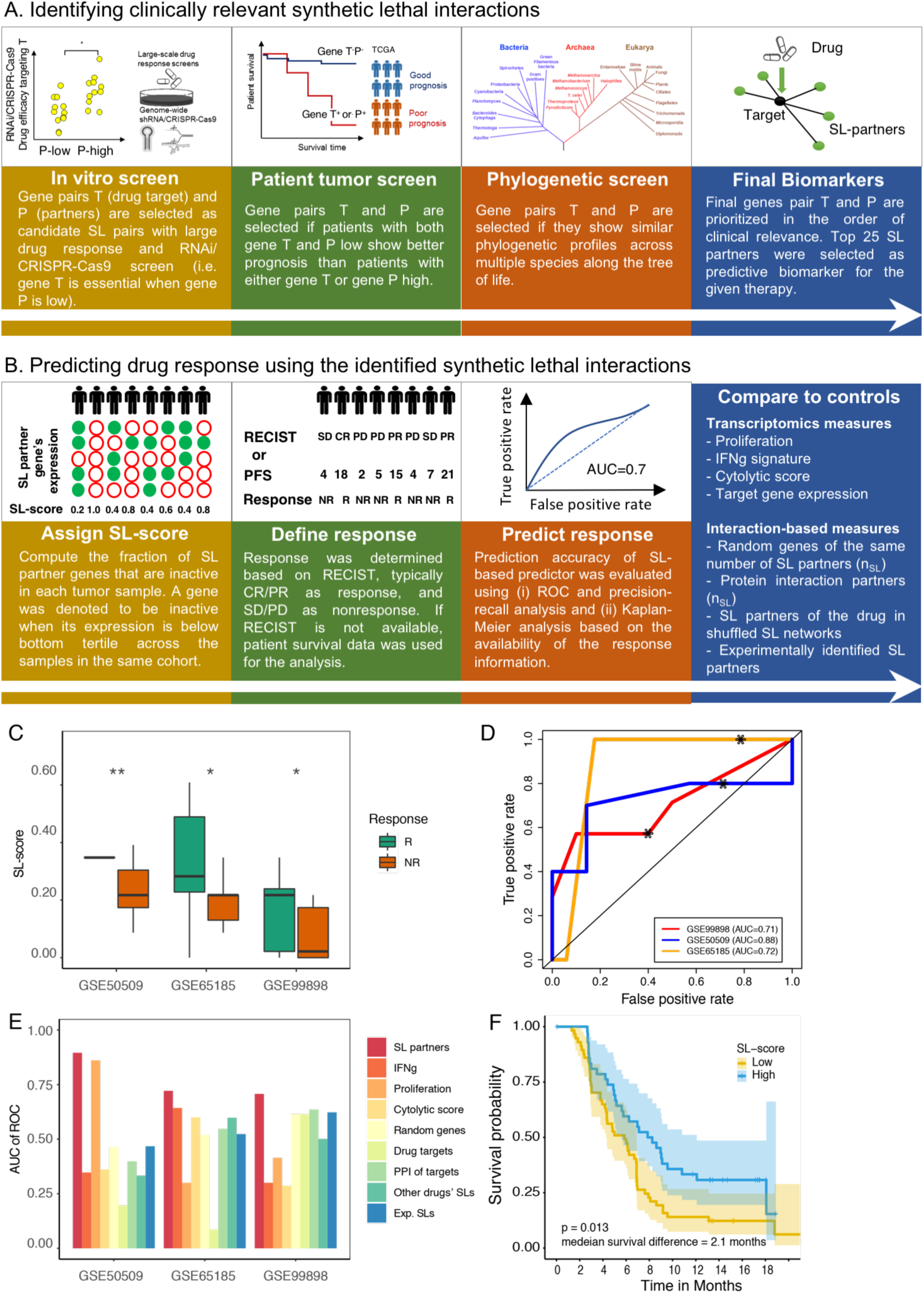
Stratifying melanoma patients for BRAF targeting treatments based on the expression of ISLE predicted BRAF SL partners. (**A**) The synthetic lethality partners (*gene P*) of the drug target genes (*gene T*) are supported by genetic dependencies in cell lines, patient tumor data, and phylogenetic profiles. (**B**) The identified SL partners of the drug target genes are used to compute an SL-score to predict the response to the given therapy. (**C**) SL-scores are significantly higher in responders (green) vs non-responders (red), based on one-sided Wilcoxon ranksum test after multiple hypothesis correction. For false discovery rates: * denotes 20% and ** denotes 10%. (**D**) ROC curves depicting the prediction accuracy of the response to BRAF inhibition using SL-scores in the three melanoma cohorts. (**E**) Bar graphs show the predictive accuracy in terms of AUC of ROC curve (Y-axis) of SL-based predictors (red) and controls including several known transcriptomics-deduced metrics (IFNg signature, proliferation index, cytolytic score, and the drug target expression levels) and several interaction-based “SL-like” scores (based on randomly chosen partners, randomly chosen PPI partners of the drug target gene(s), and the identified SL partners of other cancer drugs) in the three BRAF inhibitor cohorts (X-axis). (**F**) A Kaplan-Meier curve depicting the survival of patients with low (yellow) vs high (blue) BRAF SL-scores (dichotomized by the median SL-score). Patients with high SL-scores show better prognosis, as expected. The logrank P-value and median survival difference are denoted.

We emphasize that the treatment outcome information available in the 31 test datasets we analyzed is never used in making the response predictions. The treatment outcomes (typically provided in the form of RECIST or progression-free survival (PFS)) were only used to evaluate the resulting prediction accuracy (**Figure 1B**). Importantly, throughout the paper we use fixed sets of parameters in making predictions for targeted and immuno-therapies, respectively. Taken together, these measures are highly important as they markedly reduce the well-known risk of obtaining over-fitted predictors that fail to predict on datasets other than those on which they were originally built on (which has been observed to occur with many existing machine learning drug response predictors).

### SL-based prediction of response to targeted cancer therapies

As a proof-of-principle to assess the performance of our approach, we begin by analyzing four melanoma cohorts treated with BRAF inhibitors, where pre-treatment transcriptomics data and response information are available^14-17^. Applying the ISLE pipeline to the TCGA cohort, we identified the 25 most significant SL partners of BRAF (Methods). Reassuringly, we find that, as expected, responders have higher SL-scores than non-responders in the 3 different melanoma-BRAF cohorts for which therapy response data is available (**Figure 1C**). Quantifying the predictive power via the use of the standard area under the receiver operating characteristics curve (AUC of the ROC curve) measure, we find AUCs greater than 0.7 in all three datasets (**Figure 1D**). As some datasets do not have a balanced number of responders and nonresponders, we additionally quantified the resulting performance via precision-recall curves (often used as supplement to the routinely used ROC curves to obtain a fuller picture when evaluating prediction performance, **Figure S1A**). As evident from the latter, one can choose a single classification threshold capturing most true responders while misclassifying less than half of the non-responders. Notably, these SL based prediction accuracy levels are better overall as comparted to those obtained by several published transcriptomic based predictors, including the proliferation index^18^, IFNg signature^19^, cytolytic score^20^, or the expression of the drug target gene itself (BRAF in this case). They are also better than interaction-based scores, such as the fraction of down-regulated randomly selected genes, the fraction of *in vitro* experimentally determined SL partners, the fraction of the identified SL partners of other drugs, or the fraction of down-regulated protein-protein interaction partners (both of size similar to the SL set; empirical P<0.001) (**Figure 1E**). The fourth melanoma BRAF dataset^21^ lacks annotated RECIST response information, but we observe that patients with higher SL-scores showed significantly better treatment outcome in terms of overall survival (**Figure 1F**), as expected. BRAF’s SL partners are enriched with the functional annotation ‘regulation of GTPase mediated signal transduction’ (Fisher exact test P<0.002).

We next tested the prediction accuracy of the SL-based approach on an array of different targeted therapies and cancer types. We collected an additional cohort of 17 publicly available datasets from clinical trials of targeted therapies in cancer, each one containing both pre-treatment transcriptomics data and therapy response information. This compendium includes breast cancer patients treated with lapatinib^22^, gefinitib^23^, letrozole^24^, doxorubicin^25^, trastuzumab^26^, everolimus^27^ and cetuximab^28^; ovarian cancer patients treated with dasatinib^29^; colorectal cancer patients treated with irinotecan^30^, multiple myeloma patients treated with bortezomib^31^ and non-small cell lung cancer patients treated with sorafenib^32^. We identified the SL interaction partners of the drug targets in these datasets (Methods) and computed an SL-score for each sample using the SL partners of the corresponding drugs. Importantly, we find that higher SL-scores are associated with better response in 11 of these 15 datasets (**Figure 2A**), with AUC’s greater than 0.7 (and see precision-recall curves in **Figure S1B**), while the predictive performance of a variety of expression-based control predictors is random (**Figures 2B-C**). In the three datasets (one among the 15 datasets above and 2 additional datasets), where we have a sufficient number of samples with patient survival data, we observe that patients with higher SL-scores also have increased overall survival (**Figures 2D-F**). Taken together, these results indicate that SL interactions of the drug target genes can serve as effective biomarkers for predicting drug response across a wide range of drugs in different cancer types.

**Figure 2.**
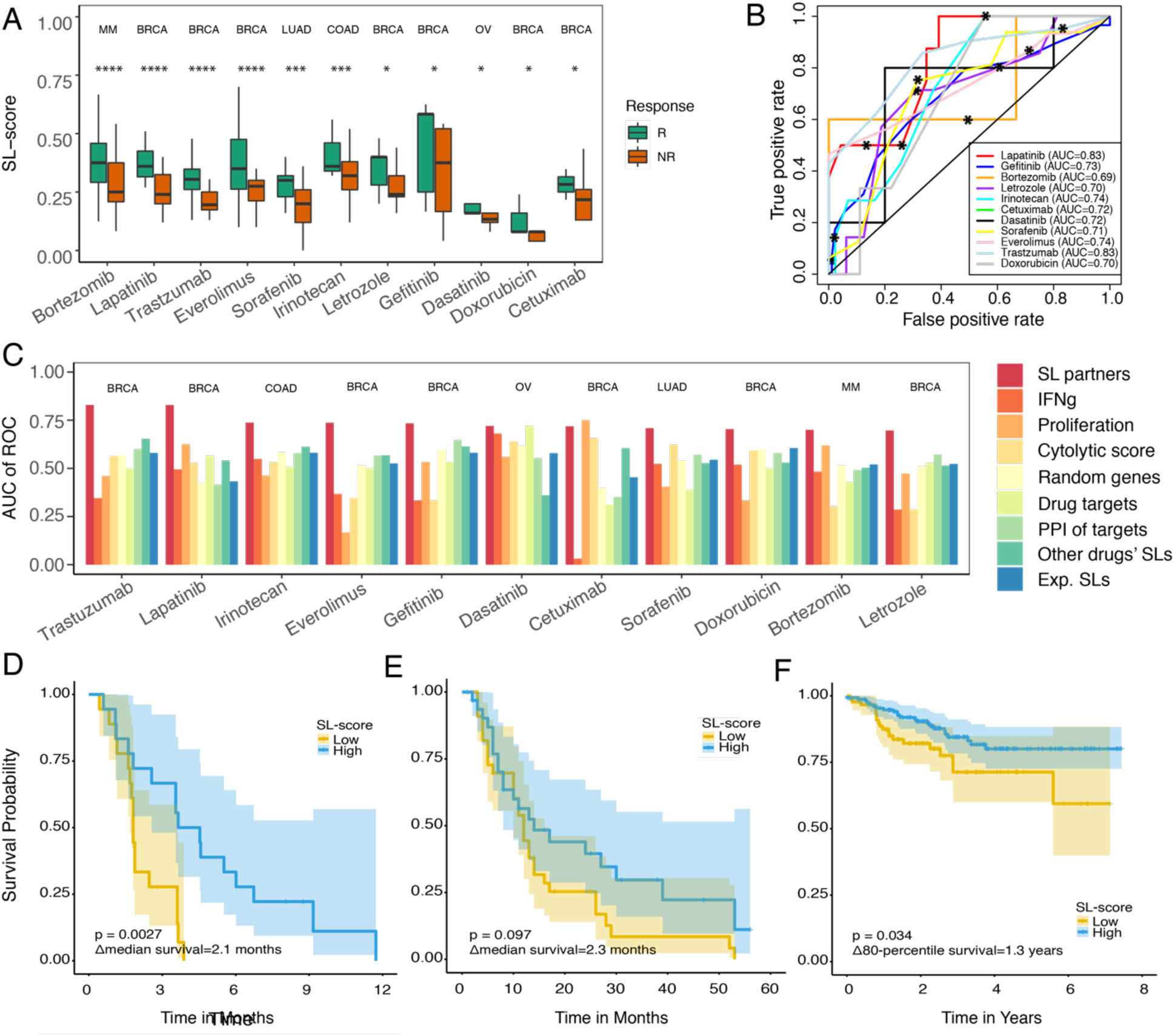
SL-based stratification of patients for targeted therapies in different cancer types. (**A**) SL-scores are significantly higher in responders (green) vs non-responders (red), based on one-sided Wilcoxon ranksum test after multiple hypothesis correction. For false discovery rates: * denotes 20%, *** denotes 5%, and **** denotes 1%. Cancer types are noted on the top of each dataset. (**B**) ROC curves for breast cancer patients treated with lapatinib (GSE66399)^22^, gefinitib (GSE33658)^23^, letrozole (GSE16391)^24^, doxorubicin (GSE8465)^25^, trastuzumab (GSE76360)^26^, everolimus (GSE119262)^27^ and cetuximab (GSE23428)^28^, ovarian cancer patients treated with dasatinib (GSE37180)^29^, non-small cell lung cancer patients treated with sorafenib (GSE33072)^32^, colorectal cancer patients treated with irinotecan (GSE72970)^30^, multiple myeloma patients treated with bortezomib (GSE68871)^31^. (**C**) Bar graphs show the predictive accuracy in terms of AUCs (Y-axis) of SL-based predictors and a variety of controls specified earlier in **Figure 1E** (X-axis). (**D-F**) Kaplan-Meier curves depicting the survival of patients with low vs high SL-scores of (**D**) Small cell lung cancer patients treated with sorafenib (GSE330712)^32^, (**E**) Colon cancer cohort treated with bevacizumab (GSE53127)^33^, and (**F**) Breast cancer cohort treated with taxane-anthracycline (GSE25055)^34^, where X-axis denotes survival time in months. Patients with high SL-scores (blue) show better prognosis than the patients with low SL-scores (yellow), as expected. The logrank P-values and median survival differences (or 80-percentile survival differences if survival exceeds 50% at the longest time point) are denoted in the figure. Tumor type abbreviations: OV, ovarian serous cystadenocarcinoma; MM, multiple myeloma; LUAD, lung adenocarcinoma; COAD, colon adenocarcinoma; BRCA, breast invasive carcinoma.

### SR-based prediction of response to checkpoint blockade

We next studied the ability of our GI-based framework to predict clinical response to checkpoint inhibitors. To this end, we introduced a few modifications into the published ISLE and INCISOR pipelines^7,8^ (Methods). We used the protein expression levels of PD1 and CTLA4 because they are likely to better reflect their activity than their mRNA levels, and we used co-culture CRISPR data to capture immune interactions modifying T-cell killing instead of the original cancer cell line screens used in the first step of ISLE and INCISOR. Using these modified versions of ISLE and INCISOR, we analyzed TCGA data to identify the SL and SR partners of PD1 and of CTLA4. We did not identify statistically significant SL interaction partners of these checkpoint genes but did find significant pan-cancer SR interactions, which were then used for further analysis. In particular, we identified two types of SR interactions, DU and DD, as we have previously defined^7,8^: SRs of the DU type denote interactions where the inactivation of the target gene is compensated by the *upregulation* of the partner rescuer gene, while SRs of the DD type denote interactions where inactivation of the target gene is compensated by the *downregulation* of the partner rescuer gene. Given a drug and a patient’s tumor expression data, we quantify the fraction *f* of DU partners that are upregulated in the tumor plus the fraction of DD partners that are downregulated. We define 1-*f* as the *SR-score*, where tumors with higher SR scores are likely to *respond* better to the given checkpoint therapy (Methods).

To evaluate the accuracy of SR based predictions, we collected a set of 11 immune checkpoint therapy datasets that included both pre-treatment transcriptomics data and therapy response information (either by RECIST or PFS). Our collection includes five melanoma datasets^35-39^ and glioma^40^, and renal cell carcinoma^41^ cohorts treated with anti-PD1 or anti-CTLA4. **Figure 3A** shows that higher SR-scores are indeed associated with better response to immune checkpoint blockade, with AUCs greater than 0.7 in 7 out of 10 datasets, where RECIST information is available (**Figure 3B**), and **Figure S2** shows the corresponding precision-recall curves. As shown in **Figure 3C**, the prediction accuracy of SR-scores is superior to a variety of expression-based controls. Notably, the SR-based framework is also predictive for a new unpublished dataset of lung adenocarcinoma (LUAD) patients treated with pembrolizumab, an anti-PD1 checkpoint inhibitor, at Samsung Medical Center (Methods, **Figure 3B-C** (denoted as ‘new SMC dataset’)). In the two datasets (including one additional dataset), where we had a sufficient patient survival data we observe that patients with higher SR-scores show significantly better prognosis for anti-PD1 therapy^39^ (**Figure 3D**) and anti-PD1/anti-CTLA4 combination^38^ (**Figure 3E**). The SR partners of PD1 and CTLA4 are enriched for IFN-gamma and antigen presentation pathways (**Figure S3**), including key immune genes such as IFNAR2, IFNGR1, and B2M, and PPI interaction partners of PD1 and CTLA4 such as B2M, RAE1, and C3. Taken together, these results testify that the SR partners of PD1 and CTLA4 serve as effective biomarkers for checkpoint response across a wide range of cancer types.

**Figure 3.**
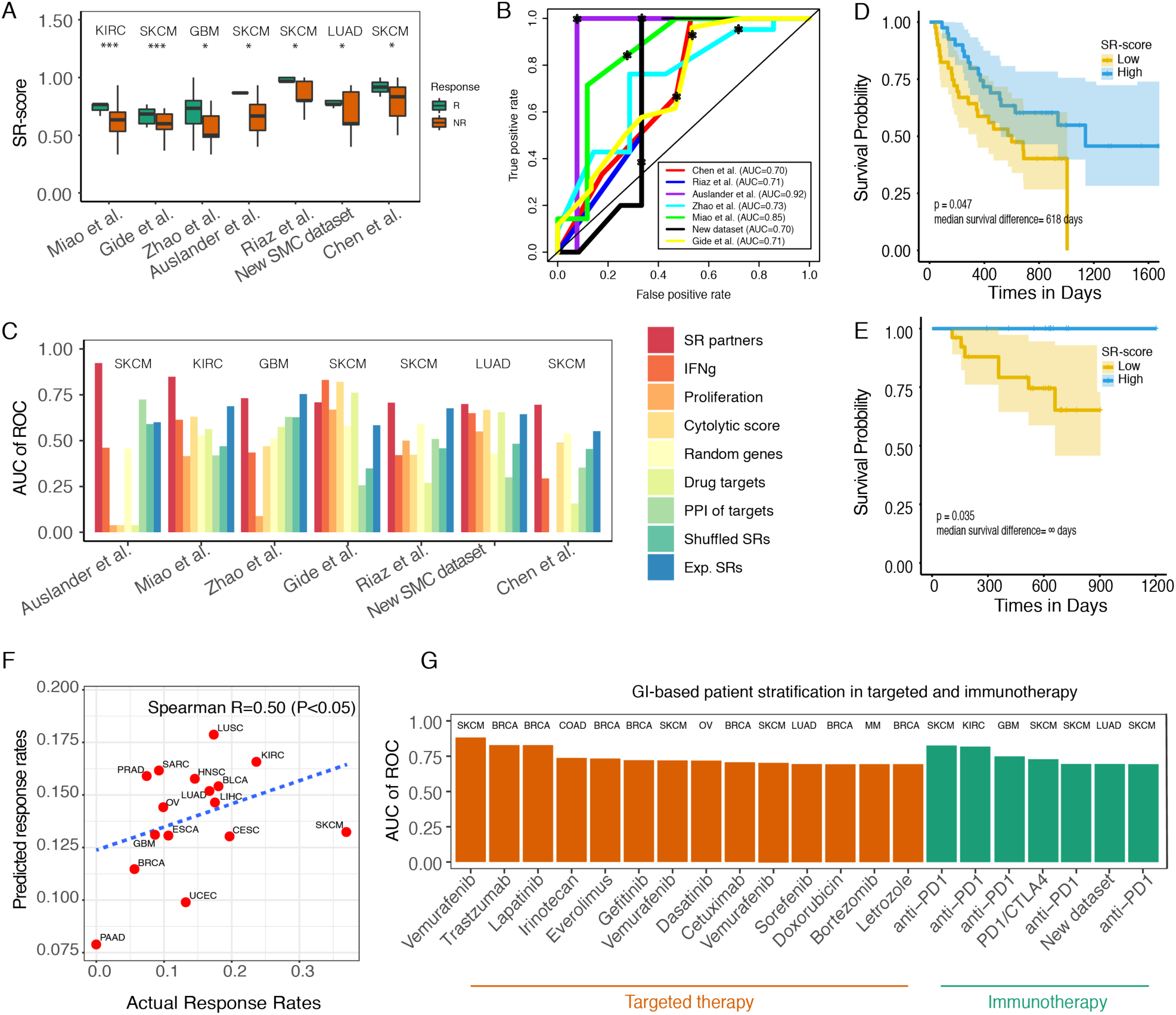
Predicting immune checkpoint therapy response based on their SR scores. (**A**) SR-scores are significantly higher in responders (green) vs nonresponders (red) based on one-sided Wilcoxon ranksum test after multiple hypothesis correction. For false discovery rates: * denotes 20%, ** denotes 10%, and *** denotes 5%. Cancer types are noted on the top of each dataset. Results are shown for melanoma cohorts^35-38^ treated with anti-PD1, anti-CTLA4 and anti-PD1 combination, renal cell carcinoma^41^, glioma^40^, and our new lung adenocarcinoma cohort treated with anti-PD1. (**B**) ROC curves showing the prediction accuracy obtained with the SR-scores across the 8 different datasets. (**C**) Bar graphs show the predictive accuracy in terms of AUC (Y-axis) of SR-based predictors and controls across different cohorts (X-axis) (control types are similar to those described in **Figure 1E**). (**D**,**E**) Kaplan-Meier curves depicting the survival of patients with low vs high anti-PD1 SR-scores in (**D**) anti-PD1-^39^ and (**E**) anti-PD1/anti-CTLA4^38^ combination-treated melanoma cohorts. Patients with high SR-scores (blue) show better prognosis than the patients with low SR-scores (yellow), and the logrank P-values and median survival differences are denoted. (**F**) The response rates predicted by the SR-scores (Y-axis) correlate with the actual response rates observed in the pertaining clinical trials (X-axis) (Spearman R=0.50, P<0.05), with a regression line (blue). Tumor type abbreviations: UCEC, uterine corpus endometrial carcinoma; SKCM, skin cutaneous melanoma; SARC, sarcoma; PRAD, prostate adenocarcinoma; PAAD, pancreatic adenocarcinoma; OV, ovarian serous cystadenocarcinoma; LUSC, lung squamous cell carcinoma; LUAD, lung adenocarcinoma; LIHC, liver hepatocellular carcinoma; KIRC, kidney renal clear cell carcinoma; HNSC, head-neck squamous cell carcinoma; GBM, glioblastoma multiforme; ESCA, esophageal carcinoma; CESC, cervical squamous cell carcinoma and endocervical adenocarcinoma; BRCA, breast invasive carcinoma; and BLCA, bladder carcinoma. **(G)** Predicting therapy response based on GI scores: The bar graphs show the overall predictive accuracy of genetic interaction-based predictors (for which we could determine the AUCs given RECIST response data, Y-axis) for targeted (red) and immunotherapies (green) in 22 different cohorts encompassing 8 cancer types and 13 treatment options (X-axis).

Next, we computed the SR scores for anti-PD1 therapy for each tumor sample in the TCGA compendium based on the expression of the SR partners (Methods). Based on the optimal classification threshold determined in the analysis presented above, we computed the fraction of responders in each cancer type and compared that to the actual response rates reported in anti-PD1 clinical trials for 16 cancer types^42^. Notably, we find that these two measures are significantly correlated (**Figure 3F**), showing that the inferred checkpoint SRs are also predictive and associated with the clinical response observed across different cancer types to these immunotherapies. Taken together, these results show that, adding to the existing determinants of response and resistance to checkpoint therapy in melanoma^43,44^, SR scores are robust predictors of response to checkpoint therapy across many different cancer types.

Summing up over all the targeted and immunotherapies we studied, our genetic interaction-based approach achieves greater than 0.7 AUC predictive performance levels in 21 out of 28 datasets containing RECIST response information, spanning 14 out of 18 targeted therapy cohorts and 7 out of 10 immunotherapy cohorts (including our new SMC dataset) (**Figure 3G**). Adding the 4 datasets where SL/SR-score is predictive of progression-free survival, our GI-based framework shows considerable predictive signals in 25 out of 32 cohorts (>78%). Notably, these accuracies are markedly better than those obtained using a range of control predictors.

Finally, we investigated if there are additional expression-based variables that could be potentially used to further refine GI based predictions in the future. Pooling together targeted therapy data to obtain a larger cohort we find that, in the patients who were predicted as responders based on their SL scores, the IFNg signature scores are significantly lower in actual non-responders vs responders (**Figure S4A**). Analogously, in a pooled analysis of checkpoint response datasets, we find that the non-responders have significantly lower IFNg and cytolytic scores than the responders (this time among the patients predicted as non-responders) (**Figure S4B**). These observations indicate that, as larger datasets are accrued, it may be worthwhile to combine GI scores and transcriptomics-based measures of immune response to enhance response prediction.

### A retrospective analysis of the WINTHER trial

To evaluate our approach in a multi-arm basket clinical trial setting we performed a retrospective analysis of the recent WINTHER trial data, the first large-scale basket clinical trial that has incorporated transcriptomics data for cancer therapy in adult patients with advanced solid tumors^1,2^. This multi-center study had two arms, one recommending treatment based on actionable mutations in a panel of cancer driver genes and another, doing so based on the patients’ transcriptomics data. We considered the gene expression data of 86 patients with 69 different targeted treatments (single or combinations) that were available. One patient had a complete response, 10 had a partial response and 12 were reported to have stable disease (labeled as responders), while 63 had a progressive disease (labeled as non-responders).

We first identified the SL partners for each of the drugs prescribed in the study. We confirmed that the resulting SL-scores of the therapies used in the trial are significantly higher in responders than non-responders (Wilcoxon ranksum P<0.05, **Figure 4A**). Notably, the SL scores of the drugs given to each patient are predictive of the actual responses observed in the trial (AUC=0.73, **Figure 4B**, with an SL-score of 0.32 as the optimal threshold that best balances precision and recall (**Figure S5**)). As shown in **Figure 4C**, the prediction accuracy of SL-score is superior to that of control expression-based predictors. This reassuring predictive signal led us to ask the following, fundamental question: how many patients would have likely benefited from the set of drugs employed in the trial, if the treatment choices for patients would have been guided by the SL-based scores of each drug in each patient, as defined and described above?

**Figure 4.**
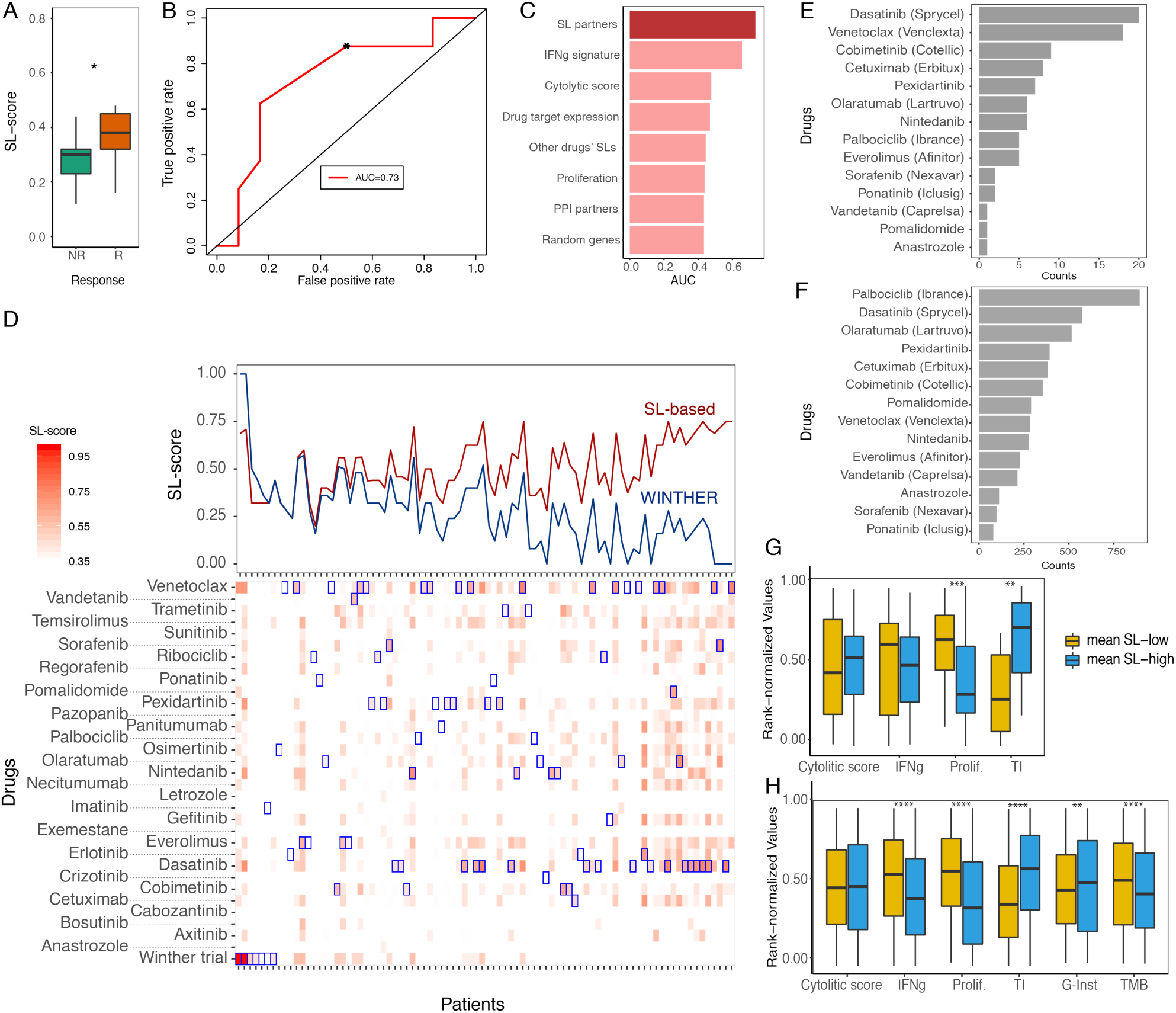
An SL-based retrospective analysis of the WINTHER trial. **(A)** Responders (CR, PR, and SD; red) show significantly higher SL-scores compared to non-responders (PD; green) (Wilcoxon ranksum P<0.01). **(B)** The ROC plot shows that the SL-scores are predictive of response to the different treatments prescribed at the trial (AUC of ROC=0.73). (**C**) Bar graphs show the predictive accuracy in terms of AUC (X-axis) of SL-based predictors and different controls (Y-axis) (control types are similar to those described in **Figure 1E**). **(D)** (**top panel**) Comparison of the SL-scores (Y-axis) of the treatments actually prescribed in the WINTHER trial (blue) and the SL-scores of the best therapy identified by our approach (red) across all 86 patients. (**bottom panel**) A more detailed display of the SL-scores (color-coded) of the treatment given in the trial (bottom row) and the SL-scores of all candidate therapies (all other rows) for all 86 patients in the trial (the treatments considered are denoted in every column). Blue boxes denote the best treatments (with highest SL-scores) recommended for each patient. **(E-F)** The bar graph shows the frequency (X-axis) of the drugs (Y-axis) predicted to be effective across patients in (**E**) the WINTHER cohort and (**F**) in a cohort of corresponding cancer types in TCGA (n=4,905). (**G-H**) The association between transcriptomics and genomics-based metrics and the mean SL-score (computed over the SL scores of all drugs considered) of each sample in (**G**) the WINTHER cohort and (**H**) in the corresponding TCGA cohort. The samples were divided into two groups, with high mean SL-score (top tertile; blue) and low mean SL-score (bottom tertile; yellow). The rank-normalized values of cytolytic activity, IFNg signature, proliferation signature, tumor mutational burden (TMB), genomic instability (G-Inst), and transcriptomic instability (TI) were compared between the two groups using a Wilcoxon ranksum test. For false discovery rates: * denotes 20%, ** denotes 10%, and *** denotes 5%, **** denotes 1%.

To answer this question in a systematic manner we considered all FDA-approved targeted anti-cancer therapies from the Drugbank database^11^ and identified the SL partners of their target genes by running the ISLE pipeline (Methods). Having predicted the SL partners for each of these drugs, we computed the SL scores of all drugs in every patient based on its tumor transcriptomics and ranked the drugs accordingly to identify the top drugs that are predicted to be most suitable for each patient. The resulting analysis shows that for more than 94% (81/86) of the patients we could identify alternate therapies that have higher SL-scores than the drugs prescribed to them in the WINTHER trial (**Figure 4D**). But given these SL-based treatment recommendations, how many may actually respond? Based on the optimal classification threshold identified earlier, 86% (74/86) of the patients are predicted to respond based on our recommendations, compared to 15% that has been reported to respond in the original trial. Out of the non-responders reported in the WINTHER trial, 87% (55/63) of the patients are predicted to be matched with effective therapies through our recommendations (**Figure 4D**). The most frequently recommended drug with the highest SL-scores in the WINTHER cohort is dasatinib, followed by venetoclax and cobimetinib (**Figure 4E**). Reassuringly, an SL-based drug coverage analysis in TCGA (**Figure 4F**), focusing on the same cancer types and drugs as those studied in the WINTHER trial, shows a similar pattern of top recommended drugs (the drug rankings show Spearman R=0.63, P<0.01). This points to the robustness of the SL-based predictions across different independent patients’ datasets.

**Figure 4D** reveals an interesting emerging pattern, where samples that display a strong SL vulnerability to one drug usually tend to have SL mediated vulnerabilities to many other drugs. We thus introduced the *mean SL-score* (computed across all drugs considered) of a given sample as a measure of its aggregate vulnerability. To investigate the molecular features that are associated with the aggregate SL vulnerability at the sample level, we considered several transcriptomics and genomics-based measures of tumor samples in both WINTHER and TCGA cohorts (**Figure 4G**,**H**). These include cytolytic activity, IFNg signature, proliferation signature, and transcriptomic instability (TI) (Methods). We additionally considered tumor mutational burden (TMB) and genomic instability (G-Inst) when analyzing the TCGA cohort, where whole exome sequencing (WES) and copy number profiles are available. Our analysis shows that transcriptomic and genomic instabilities are significantly higher in the samples with high mean SL-score in both cohorts. Notably, this finding indicates that SL-based treatment opportunities may actually increase in advanced tumors, which have increases G-Inst and TI levels.

## Discussion

We have demonstrated that by mining large-scale “-omics” data from patients’ tumors one can computationally infer candidate pairs of genetic interactions that can be used as predictive companion diagnostics for many targeted and immunotherapy treatments, across multiple cancer types. The resulting prediction accuracy is of potential translational value for many of the drugs tested. Furthermore, as shown in the analysis of the WINTHER trial, its application may significantly increase the number of patients that can benefit from precision-based treatments.

Among the drugs that were commonly predicted to be of clinical utility in our analysis of the latter trial, dasatinib and venetoclax are approved for hematopoietic cancers but their potential use for solid tumors has been investigated in multiple solid tumors including colorectal and breast cancer^45,46^. Cobimetinib is approved for the treatment of melanoma, and clinical trials are ongoing for multiple tumors including craniopharyngioma, non-small cell lung cancer, pancreatic cancer, and ovarian cancer. Notably, olaratumab, ranked 6^th^ in our analysis, is a monoclonal antibody developed for the treatment of solid tumors, directed against the platelet-derived growth factor receptor alpha (PDGFRA)^47^. However, olaratumab has been removed from the US and the European markets due to insufficient proof of efficacy. These drugs also attain wide coverage in the independent analysis of the TCGA cohort. Taken together, these results lay a basis for future prospective studies based on SL-based stratification of patients’ treatments from their tumor transcriptomics.

Our predictor is fundamentally different from previous efforts for therapy response prediction in several important ways: (1) The GIs underlying the prediction are inferred from analyzing pre-treatment data from the TCGA ensemble without considering any treatment labels; thus, the resulting predictors are likely to be less prone to the risk of overfitting arising when applying machine learning to build predictors based on the relatively small training datasets that are currently available to us. Furthermore, the GIs used in this study are inferred from a pan cancer analysis of the TCGA – that is, they are common across many different cancer types; as such, they are less context-sensitive and more likely to be predictive in different cancer types. (2) The interactions enabling the predictions have clear biological meanings as functional GIs and their scoring is simple and intuitive, differing from the typical “black-box” solutions characteristic of machine learning approaches. (3) Finally, the resulting predictive performance is far superior to alternative predictors. As far as we are aware, no single model has previously been shown to obtain such predictive accuracies across so many targeted and immunotherapy datasets.

Importantly, we anticipate that the current levels of predictive performance can and will be further improved in the foreseen future. This will likely be achieved by analyzing even larger patient cohorts of different cancer types as those accumulate, by considering the accumulating tumor proteomic data and by developing computational approaches for inferring genetic interactions from single cell sequencing data, which will also enable one to identify genetic interactions that are grounded on specific cell types (e.g., between tumor cells and CD8^+^ T cell in their environment). As more data accumulates, we may learn how to combine SL and SR interactions together to further boost prediction performance. Finally, the GI-based stratification signatures of each drug are biologically interpretable and in many cases are rather small, so they can be measured in future trials in a cost-effective manner.

In summary, our work presents a new paradigm harnessing genetic interaction networks to advance precision cancer medicine, by systematically analyzing patients’ tumor transcriptomics data. Our results show that computationally inferred genetic interaction partners of the drug targets are predictive of clinical outcomes of cancer treatment when applied to a large collection of patient transcriptomics datasets, without any training on the latter. Combined with the present precision oncology approaches, the genetic interaction-based strategy presented here may provide a transformative way to further advance precision oncology, motivating prospective transcriptomics-based clinical trials to further test its value for treating patients with advanced malignancies.

## Methods

### Data collection

We collected the patients’ pre-treatment gene expression data with therapy response information from public databases. Our search was focused on GEO and ArrayExpress but also include general literature search using different combinations of the following search terms: ‘drug response patient cancer pre-treatment expression transcriptomics therapy resistance signature’ in March 2019. We charted 21 targeted (drugs of specific targets) and 10 immune checkpoint therapies covering a total of 10 cancer types treated with 15 drugs. In addition, we analyzed a new unpublished dataset of lung adenocarcinoma patients treated with pembrolizumab at Samsung Medical Center.

### Identifying genetic interaction networks

To identify clinically relevant SL interactions for targeted therapies, we followed the three-step procedure described in ^7^. We first created initial pool of SL pairs identified in cell lines via RNAi/CRISPR-Cas9^12,13^ or pharmacological screens^48,49^. Second, among the candidate gene pairs from the first step, we selected those gene pairs whose co-inactivation is associated with better prognosis in patients, testifying that they may hamper tumor progression. Third, we prioritized SL paired genes with similar phylogenetic profiles across different species. The top 25 clinically relevant partners were selected among the significant partners for each drug. For the cases where a drug has multiple targets, we selected the top partners that are significant among all SL partners of the target genes of the drug. We have confirmed that the predictive signal does not sensitively depend on the number of the top SL partners.

To identify GIs for immunotherapy, we introduced a few modifications to our general GI inference pipeline previously introduced^7,8^: (1) We used the protein expression levels of PD1 and CTLA4, as these are more likely to better represent the protein activity than its mRNA levels, and this data for PD1 and CTLA4 is available in TCGA (as reverse phase protein lysate microarray (RPPA) values). We considered the level of expression of PD1 (or CTLA4) on CD8^+^ T cells, which is computed as the ratio between PD1 (or CTLA4) RPPA values and the computationally estimated CD8^+^ T-cell abundance^50^. For the GI partner levels, we resorted to their gene expression and somatic copy number alterations (SCNA) data as referenced in previous studies^7,8^ because protein expression was measured only for a small subset of genes. (2) Instead of using the genome-scale functional screens in cancer cell lines (DepMap^12,13^) in the first step of ISLE or INCISOR, we analyze genome-wide CRISPR screens performed in co-cultures of cancer and T-cells, identifying genes whose knock-out enhances or prevents T-cell mediated killing. To this end we incorporated four such genome-wide CRISPR screens publicly available^51-54^. Since two of them involve treatment of anti-PD1/PDL1, we used these two as the basis for identifying GIs involved in anti-PD1 response prediction^52,54^; whereas the remaining two screens, identifying those genes involved in generic immune response^51,53^, were used to infer GIs for anti-CTLA4 response prediction. (3) We focused on the mediators of resistance to immune checkpoint therapies using synthetic rescue (SR) interactions, as no statistically significant SL interaction partners were identified via ISLE for either PD1 or CTLA4. The top clinically relevant partners for each of DU and DD type SR interactions were selected among the significant partners for immune-checkpoint predictions. We have confirmed that the predictive signal does not sensitively depend on the number of the top SR partners.

We analyzed TCGA data applying this immune-adapted INCISOR pipeline to identify pan cancer SR interactions that are more likely to be clinically relevant across many cancer types. In particular, we considered two types of SR interactions, defined and studied first in ^8^, DU and DD: SRs of the DU type denote interactions where inactivation of the target gene is compensated by the *upregulation* of the partner rescuer gene, while SRs of the DD type denote interactions where inactivation of the target gene is compensated by the *downregulation* of the partner rescuer gene. For the melanoma^37^ cohort where only a subset of genes’ expression were measured by nanostring, we performed our analysis only for that subset of genes available in the specific cohorts.

### Predicting drug response in patients using GI partners

We used the identified SL or SR partners for drug response prediction. We define SL-score for targeted therapy, that is the fraction of the number of inactive SL partners in a given sample multiplied by an indicator variable that denotes whether the target gene expression is within 70-percentile in the given sample. We used SR-score to predict response to immunotherapy, that quantifies the fraction of overactive DU-type SR partners and inactive DD-type SR partners. In particular, we calculate SR-score, that quantifies the fraction *f*, of DU partners that are active and the number of DD partners that are inactive, and we used 1-*f* as the SR-score to predict *responders*. Higher SL- or SR-score is predictive of response to therapies. In each patient drug response dataset, a gene is determined to be inactive (or overactive) if its expression is below bottom tertile (or above top tertile) across samples in the same dataset following the previous studies^7,8^. Using the computed SL/SR-scores, we solved either classification problem to predict responders or performed Kaplan-Meier analysis for predicting patient survival, depending on the availability of the data. For the datasets where the response information is available in the form of RECIST criteria, we solved classification problem. For the cases where progression-free survival time is available for all patients (with no censoring event), we used the median progression-free survival from the relevant literature as the cutoff to distinguish the responders from nonresponders and solved a classification problem. For the datasets where we only have overall or progression-free survival with censoring information, we performed Kaplan-Meier analysis.

### TCGA anti-PD1 coverage analysis

The objective response rates of anti-PD1 therapy in each cancer type in TCGA were predicted via our SR interaction partners of PD1 identified above. We computed the SR scores in each tumor sample in the TCGA compendium, based on its transcriptomics profiles following the above definition of SR-score, and labeled it as responder or non-responder accordingly using precision-recall break-even point as threshold across all 8 immune checkpoint datasets, where our SR-score is predictive. Using this fixed cut-off, we then computed the fractions of responders for each cancer type and compared these with the actual response rates reported in anti-PD1 clinical trials for 16 cancer types where the data is available^42^ using Spearman rank correlation. In each patient drug response datasets, a gene is determined to be inactive (or overactive) if its expression is below bottom tertile (or above top tertile) across samples in the same dataset following the previous studies^7,8^.

### Retrospective analysis of WINTHER trial

The trial involved 10 different cancer types, mostly colon, lung and head and neck cancer. This trial has two arms, one recommending treatment based on actionable mutations in a panel of cancer driver genes and another based on the patients’ transcriptomics data. We considered the Agilent microarray data of 86 patients with 69 different targeted treatments (single or combinations) that were available to us. One patient had a complete response, 10 had a partial response and 12 were reported to have stable disease (labeled as responders in our analysis), while 63 had a progressive disease (labeled as non-responders). For each of the drug that was used in the trial, we identified their target genes’ SL partners following our computational pipeline described above. We calculated SL-score of the therapy used in the WINTHER trial and quantify the accuracy of SL-score in predicting response across the 86 patients via ROC analysis. We focused on the patients who received monotherapy to compute the AUC of ROC curves, since our subsequent analysis is focused on monotherapies. To determine alternative treatment options recommended by SL-based approach, we then considered all FDA-approved targeted therapies obtained from the Drugbank database and the literature as candidates. We then identified SL partners of the drug target genes, where we find significant SL partners for each drug. Having predicted these SL partners for each of these drugs, we finally computed SL-score of those drugs for all patients using the identified SL-network and the patients’ transcriptomic data and ranked the drugs accordingly to identify the top drugs that are predicted to be most suitable for each patient. To evaluate the robustness of our approach, we repeated the same analysis using the matching cancer types in TCGA cohorts. We define the SL vulnerability as the mean SL-score of a given tumor among all the SL-scores of different drugs. We checked the association between different tumor molecular phenotypes and the SL vulnerability, where the tumor samples were divided into two groups with high SL vulnerability (top tertile) and low SL vulnerability (bottom tertile), and Wilcoxon ranksum test was performed for the comparison. We considered the following phenotypes: cytolytic activity, IFNg signature, proliferation signature, transcriptomic instability in WINTHER and TCGA cohorts. Transcriptomic instability quantifies the genome-wide extent of the transcriptomic dysregulation of genes by computing the fraction of over- or under-expressed genes among all protein coding genes. The over- (or under-) expression of a gene is defined as the top or bottom tertile of the expression of the given gene in the reference population. If we have sufficient number of samples with the same cancer type in the same dataset, those were used as reference; if we do not have sufficient number of samples with the same cancer type in the same dataset, we used TCGA samples of the same cancer type as reference. Genomic instability and tumor mutational burden were considered only in TCGA where whole exome sequencing and copy number profiles are available.

## Acknowledgements

This research is supported in part by the Intramural Research Program of the National Institutes of Health (NIH), National Cancer Institute (NCI), Center for Cancer Research (CCR). This work utilized the computational resources of the NIH HPC Biowulf cluster. We thank Peng Jiang, Alejandro A. Schäffer, E. Michael Gertz, and Sanna Madan at Cancer Data Science Lab, CCR/NCI/NIH for insightful comments.

## References

1 Rodon, J. et al. Genomic and transcriptomic profiling expands precision cancer medicine: the WINTHER trial. Nat Med 25, 751–758, doi:10.1038/s41591-019-0424-4 (2019).

2 Beaubier, N. et al. Integrated genomic profiling expands clinical options for patients with cancer. Nat Biotechnol, doi:10.1038/s41587-019-0259-z (2019).

3 Vaske, O. M. et al. Comparative Tumor RNA Sequencing Analysis for Difficult-to-Treat Pediatric and Young Adult Patients With Cancer. JAMA Netw Open 2, e1913968, doi:10.1001/jamanetworkopen.2019.13968 (2019).

4 Tanioka, M. et al. Integrated Analysis of RNA and DNA from the Phase III Trial CALGB 40601 Identifies Predictors of Response to Trastuzumab-Based Neoadjuvant Chemotherapy in HER2-Positive Breast Cancer. Clin Cancer Res 24, 5292–5304, doi:10.1158/1078-0432.CCR-17-3431 (2018).

5 Lord, C. J. & Ashworth, A. PARP inhibitors: Synthetic lethality in the clinic. Science 355, 1152–1158, doi:10.1126/science.aam7344 (2017).

6 Eischen, C. M., Woo, D., Roussel, M. F. & Cleveland, J. L. Apoptosis triggered by Myc-induced suppression of Bcl-X(L) or Bcl-2 is bypassed during lymphomagenesis. Mol Cell Biol 21, 5063–5070, doi:10.1128/MCB.21.15.5063-5070.2001 (2001).

7 Lee, J. S. et al. Harnessing synthetic lethality to predict the response to cancer treatment. Nat Commun 9, 2546, doi:10.1038/s41467-018-04647-1 (2018).

8 Sahu, A. D. et al. Genome-wide prediction of synthetic rescue mediators of resistance to targeted and immunotherapy. Mol Syst Biol 15, e8323, doi:10.15252/msb.20188323 (2019).

9 Feng, X. et al. A Platform of Synthetic Lethal Gene Interaction Networks Reveals that the GNAQ Uveal Melanoma Oncogene Controls the Hippo Pathway through FAK. Cancer Cell 35, 457–472 e455, doi:10.1016/j.ccell.2019.01.009 (2019).

10 Pathria, G. et al. Translational reprogramming marks adaptation to asparagine restriction in cancer. Nat Cell Biol, doi:10.1038/s41556-019-0415-1 (2019).

11 Law, V. et al. DrugBank 4.0: shedding new light on drug metabolism. Nucleic Acids Res 42, D1091–1097, doi:10.1093/nar/gkt1068 (2014).

12 Tsherniak, A. et al. Defining a Cancer Dependency Map. Cell 170, 564–576 e516, doi:10.1016/j.cell.2017.06.010 (2017).

13 Meyers, R. M. et al. Computational correction of copy number effect improves specificity of CRISPR-Cas9 essentiality screens in cancer cells. Nat Genet 49, 1779–1784, doi:10.1038/ng.3984 (2017).

14 Kakavand, H. et al. PD-L1 Expression and Immune Escape in Melanoma Resistance to MAPK Inhibitors. Clin Cancer Res 23, 6054–6061, doi:10.1158/1078-0432.CCR-16-1688 (2017).

15 Rizos, H. et al. BRAF inhibitor resistance mechanisms in metastatic melanoma: spectrum and clinical impact. Clin Cancer Res 20, 1965–1977, doi:10.1158/1078-0432.CCR-13-3122 (2014).

16 Hugo, W. et al. Non-genomic and Immune Evolution of Melanoma Acquiring MAPKi Resistance. Cell 162, 1271–1285, doi:10.1016/j.cell.2015.07.061 (2015).

17 McArthur, G. A. et al. Safety and efficacy of vemurafenib in BRAF(V600E) and BRAF(V600K) mutation-positive melanoma (BRIM-3): extended follow-up of a phase 3, randomised, open-label study. Lancet Oncol 15, 323–332, doi:10.1016/S1470-2045(14)70012-9 (2014).

18 Whitfield, M. L., George, L. K., Grant, G. D. & Perou, C. M. Common markers of proliferation. Nat Rev Cancer 6, 99–106, doi:10.1038/nrc1802 (2006).

19 Ayers, M. et al. IFN-gamma-related mRNA profile predicts clinical response to PD-1 blockade. J Clin Invest 127, 2930–2940, doi:10.1172/JCI91190 (2017).

20 Rooney, M. S., Shukla, S. A., Wu, C. J., Getz, G. & Hacohen, N. Molecular and genetic properties of tumors associated with local immune cytolytic activity. Cell 160, 48–61, doi:10.1016/j.cell.2014.12.033 (2015).

21 Wongchenko, M. J. et al. Gene Expression Profiling in BRAF-Mutated Melanoma Reveals Patient Subgroups with Poor Outcomes to Vemurafenib That May Be Overcome by Cobimetinib Plus Vemurafenib. Clin Cancer Res 23, 5238–5245, doi:10.1158/1078-0432.CCR-17-0172 (2017).

22 Guarneri, V. et al. Prospective Biomarker Analysis of the Randomized CHER-LOB Study Evaluating the Dual Anti-HER2 Treatment With Trastuzumab and Lapatinib Plus Chemotherapy as Neoadjuvant Therapy for HER2-Positive Breast Cancer. Oncologist 20, 1001–1010, doi:10.1634/theoncologist.2015-0138 (2015).

23 Massarweh, S. et al. A phase II neoadjuvant trial of anastrozole, fulvestrant, and gefitinib in patients with newly diagnosed estrogen receptor positive breast cancer. Breast Cancer Res Treat 129, 819–827, doi:10.1007/s10549-011-1679-8 (2011).

24 Desmedt, C. et al. The Gene expression Grade Index: a potential predictor of relapse for endocrine-treated breast cancer patients in the BIG 1-98 trial. BMC Med Genomics 2, 40, doi:10.1186/1755-8794-2-40 (2009).

25 Julka, P. K. et al. A phase II study of sequential neoadjuvant gemcitabine plus doxorubicin followed by gemcitabine plus cisplatin in patients with operable breast cancer: prediction of response using molecular profiling. Br J Cancer 98, 1327–1335, doi:10.1038/sj.bjc.6604322 (2008).

26 Varadan, V. et al. Immune Signatures Following Single Dose Trastuzumab Predict Pathologic Response to PreoperativeTrastuzumab and Chemotherapy in HER2-Positive Early Breast Cancer. Clin Cancer Res 22, 3249–3259, doi:10.1158/1078-0432.CCR-15-2021 (2016).

27 Sabine, V. S. et al. Gene expression profiling of response to mTOR inhibitor everolimus in pre-operatively treated post-menopausal women with oestrogen receptor-positive breast cancer. Breast Cancer Res Treat 122, 419–428, doi:10.1007/s10549-010-0928-6 (2010).

28 Carey, L. A. et al. TBCRC 001: randomized phase II study of cetuximab in combination with carboplatin in stage IV triple-negative breast cancer. J Clin Oncol 30, 2615–2623, doi:10.1200/JCO.2010.34.5579 (2012).

29 Secord, A. A. et al. A phase I trial of dasatinib, an SRC-family kinase inhibitor, in combination with paclitaxel and carboplatin in patients with advanced or recurrent ovarian cancer. Clin Cancer Res 18, 5489–5498, doi:10.1158/1078-0432.CCR-12-0507 (2012).

30 Del Rio, M. et al. Molecular subtypes of metastatic colorectal cancer are associated with patient response to irinotecan-based therapies. Eur J Cancer 76, 68–75, doi:10.1016/j.ejca.2017.02.003 (2017).

31 Terragna, C. et al. The genetic and genomic background of multiple myeloma patients achieving complete response after induction therapy with bortezomib, thalidomide and dexamethasone (VTD). Oncotarget 7, 9666–9679, doi:10.18632/oncotarget.5718 (2016).

32 Byers, L. A. et al. An Epithelial-Mesenchymal Transition Gene Signature Predicts Resistance to EGFR and PI3K Inhibitors and Identifies Axl as a Therapeutic Target for Overcoming EGFR Inhibitor Resistance. Clinical Cancer Research 19, 279–290, doi:10.1158/1078-0432.Ccr-12-1558 (2013).

33 Pentheroudakis, G. et al. A study of gene expression markers for predictive significance for bevacizumab benefit in patients with metastatic colon cancer: a translational research study of the Hellenic Cooperative Oncology Group (HeCOG). BMC Cancer 14, 111, doi:10.1186/1471-2407-14-111 (2014).

34 Hatzis, C. et al. A Genomic Predictor of Response and Survival Following Taxane-Anthracycline Chemotherapy for Invasive Breast Cancer. Jama-Journal of the American Medical Association 305, 1873–1881, doi:10.1001/jama.2011.593 (2011).

35 Riaz, N. et al. Tumor and Microenvironment Evolution during Immunotherapy with Nivolumab. Cell 171, 934–949 e915, doi:10.1016/j.cell.2017.09.028 (2017).

36 Auslander, N. et al. Robust prediction of response to immune checkpoint blockade therapy in metastatic melanoma. Nat Med 24, 1545–1549, doi:10.1038/s41591-018-0157-9 (2018).

37 Chen, P. L. et al. Analysis of Immune Signatures in Longitudinal Tumor Samples Yields Insight into Biomarkers of Response and Mechanisms of Resistance to Immune Checkpoint Blockade. Cancer Discov 6, 827–837, doi:10.1158/2159-8290.CD-15-1545 (2016).

38 Gide, T. N., Wilmott, J. S., Scolyer, R. A. & Long, G. V. Primary and Acquired Resistance to Immune Checkpoint Inhibitors in Metastatic Melanoma. Clin Cancer Res 24, 1260–1270, doi:10.1158/1078-0432.CCR-17-2267 (2018).

39 Liu, D. et al. Integrative molecular and clinical modeling of clinical outcomes to PD1 blockade in patients with metastatic melanoma. Nat Med 25, 1916–1927, doi:10.1038/s41591-019-0654-5 (2019).

40 Zhao, J. et al. Immune and genomic correlates of response to anti-PD-1 immunotherapy in glioblastoma. Nat Med 25, 462–469, doi:10.1038/s41591-019-0349-y (2019).

41 Miao, D. et al. Genomic correlates of response to immune checkpoint therapies in clear cell renal cell carcinoma. Science 359, 801–806, doi:10.1126/science.aan5951 (2018).

42 Yarchoan, M., Hopkins, A. & Jaffee, E. M. Tumor Mutational Burden and Response Rate to PD-1 Inhibition. N Engl J Med 377, 2500–2501, doi:10.1056/NEJMc1713444 (2017).

43 Keenan, T. E., Burke, K. P. & Van Allen, E. M. Genomic correlates of response to immune checkpoint blockade. Nat Med 25, 389–402, doi:10.1038/s41591-019-0382-x (2019).

44 Fares, C. M., Van Allen, E. M., Drake, C. G., Allison, J. P. & Hu-Lieskovan, S. Mechanisms of Resistance to Immune Checkpoint Blockade: Why Does Checkpoint Inhibitor Immunotherapy Not Work for All Patients? Am Soc Clin Oncol Educ Book 39, 147–164, doi:10.1200/EDBK_240837 (2019).

45 Lok, S. W. et al. A Phase Ib Dose-Escalation and Expansion Study of the BCL2 Inhibitor Venetoclax Combined with Tamoxifen in ER and BCL2-Positive Metastatic Breast Cancer. Cancer Discov 9, 354–369, doi:10.1158/2159-8290.CD-18-1151 (2019).

46 Scott, A. J. et al. Evaluation of the efficacy of dasatinib, a Src/Abl inhibitor, in colorectal cancer cell lines and explant mouse model. PLoS One 12, e0187173, doi:10.1371/journal.pone.0187173 (2017).

47 Tap, W. D. et al. Olaratumab and doxorubicin versus doxorubicin alone for treatment of soft-tissue sarcoma: an open-label phase 1b and randomised phase 2 trial. Lancet 388, 488–497, doi:10.1016/S0140-6736(16)30587-6 (2016).

48 Basu, A. et al. An Interactive Resource to Identify Cancer Genetic and Lineage Dependencies Targeted by Small Molecules. Cell 154, 1151–1161, doi:10.1016/j.cell.2013.08.003 (2013).

49 Iorio, F. et al. A Landscape of Pharmacogenomic Interactions in Cancer. Cell 166, 740–754, doi:10.1016/j.cell.2016.06.017 (2016).

50 Newman, A. M. et al. Robust enumeration of cell subsets from tissue expression profiles. Nat Methods 12, 453–457, doi:10.1038/nmeth.3337 (2015).

51 Patel, S. J. et al. Identification of essential genes for cancer immunotherapy. Nature 548, 537–542, doi:10.1038/nature23477 (2017).

52 Manguso, R. T. et al. In vivo CRISPR screening identifies Ptpn2 as a cancer immunotherapy target. Nature 547, 413–418, doi:10.1038/nature23270 (2017).

53 Pan, D. et al. A major chromatin regulator determines resistance of tumor cells to T cell-mediated killing. Science 359, 770–775, doi:10.1126/science.aao1710 (2018).

54 Kearney, C. J. et al. Tumor immune evasion arises through loss of TNF sensitivity. Sci Immunol 3, doi:10.1126/sciimmunol.aar3451 (2018).

