## Supplementary Information for "Whole exome precision oncology targeting synthetic lethal vulnerabilities across the tumor transcriptome"

### 1. Supplementary Figures

**A** **B**

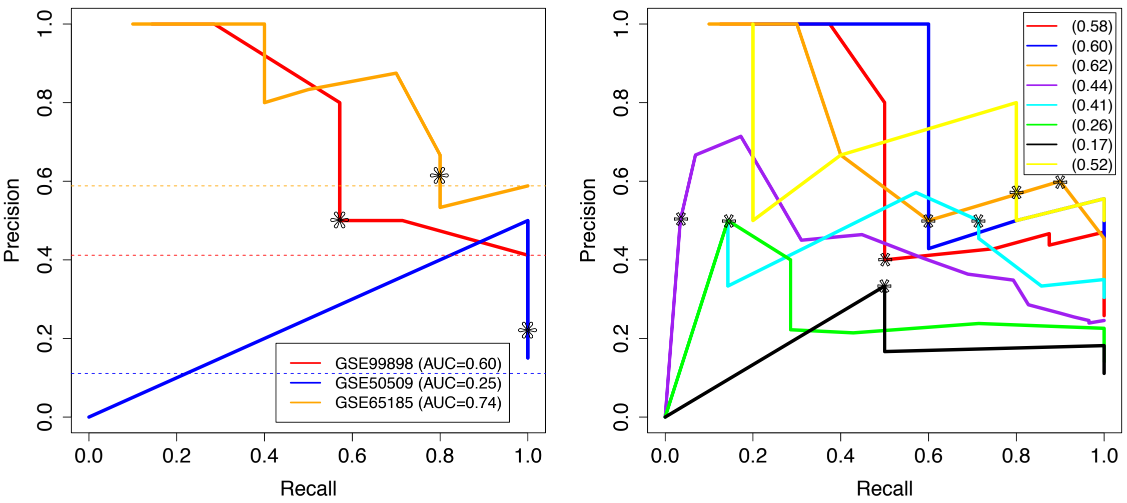


**Figure S1. Stratifying patients for targeted therapies based on the expression of ISLE predicted SL partners.** (**A**) Precision-recall curves depicting the prediction accuracy of the response to BRAF inhibition using SL-scores in the three melanoma cohorts. (**B**) Precision-recall curves for breast cancer patients treated with lapatinib (red; GSE66399)[1], gefinitib (blue; GSE33658)[2], letrozole (purple; GSE16391)[3], doxorubicin (grey; GSE8465)[4], trastuzumab (light blue; GSE76360)[5], everolimus (pink; GSE119262)[6] and cetuximab (green; GSE23428)[7], ovarian cancer patients treated with dasatinib (black; GSE37180)[8], non-small cell lung cancer patients treated with sorafenib (yellow; GSE33072)[9], colorectal cancer patients treated with irinotecan (cyan; GSE72970)[10], multiple myeloma patients treated with bortezomib (orange; GSE68871)[11]. The AUC of each precision-recall curve is denoted in the figure legend.

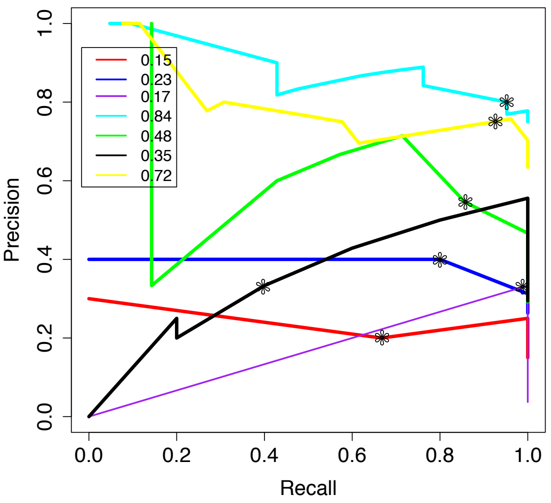


**Figure S2. Stratifying patients for immunotherapies based on the expression of NCISOR predicted SR partners.** Precision-recall curves for Chen et al. (red)[12], Riaz et al. (blue)[13], Auslander et al. (purple)[14], Zhao et al. (cyan)[15], Miao et al. (green)[16], New SMC dataset (black), and Gide et al. (yellow)[17]. The AUC of each precision-recall curve is denoted in the figure legend.

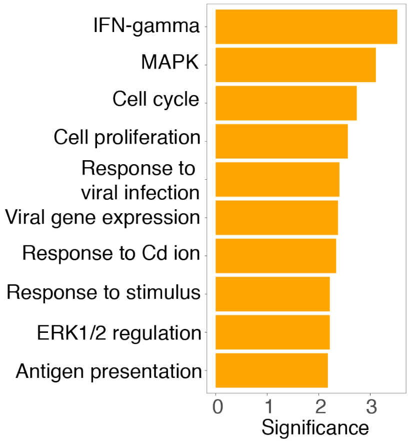


**Figure S3.** Pathway enrichment of the predicted SR partners (Y-axis) and their hypergeometric enrichment levels (X-axis).

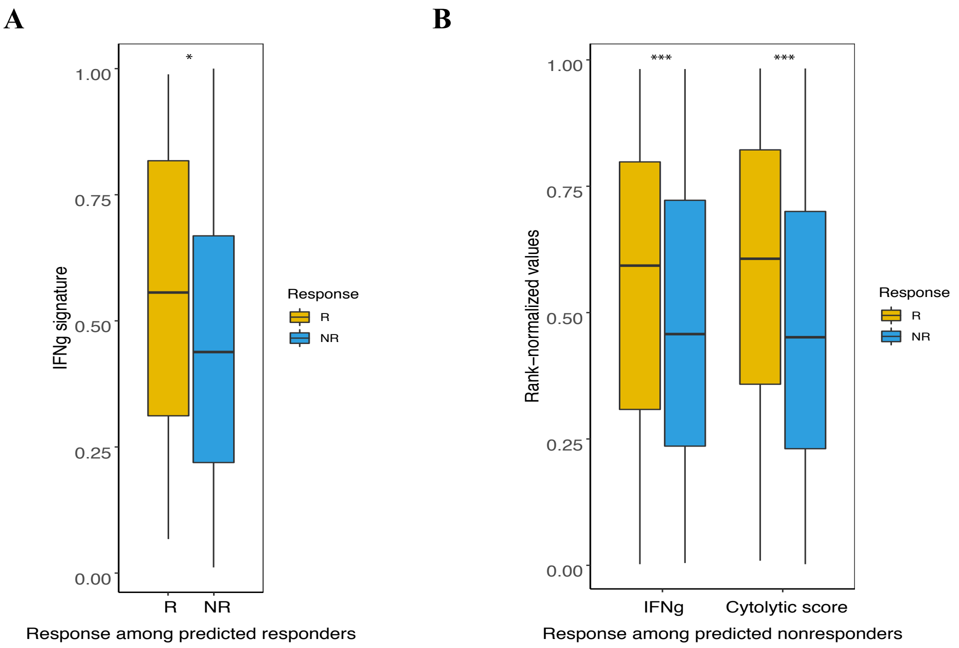


**Figure S4. Identifying additional factors that predict response. (A)** Boxplots showing the significant differences in IFNg signature levels between the responders (yellow) and nonresponders (blue), among the patients predicted as responders to targeted therapy based on SL-score. **(B)** Similarly, for SR-score predicted non-responders for immune checkpoint therapy.

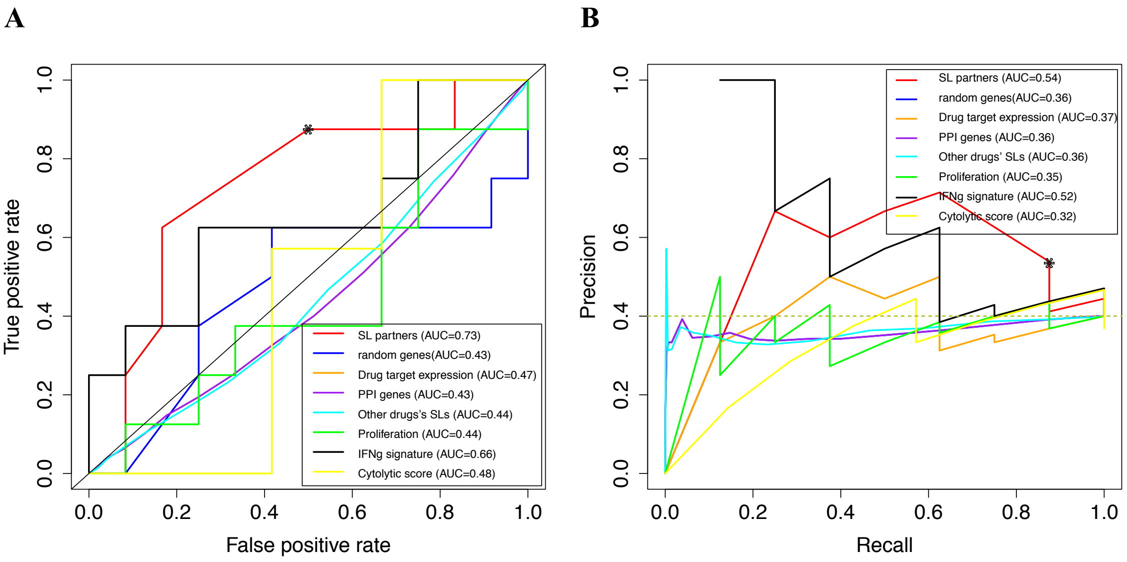


**Figure S5. Predicting therapy response across different drugs and tumor types in WINTHER trial data.** (**A**) The ROC plot shows that the SL-scores are predictive of response to the different treatments prescribed at the trial (AUC of ROC=0.73), which is higher than those control predictors that are similar to those described in **Figure 1E**). (**B**) The precision-recall curve shows that the SL-scores are predictive of response to the different treatments prescribed at the trial (AUC of precision-recall curve=0.54), which is higher than those control predictors that are similar to those described in **Figure 1E**).
